# Planning a future randomized clinical trial based on a network of relevant past trials

**DOI:** 10.1101/324871

**Authors:** Georgia Salanti, Adriani Nikolakopoulou, Alex J Sutton, Stephan Reichenbach, Sven Trelle, Huseyin Naci, Matthias Egger

## Abstract

*Background:* The important role of network meta-analysis of randomized clinical trials in health technology assessment and guideline development is increasingly recognized. This approach has the potential to obtain conclusive results earlier than with new standalone trials or conventional, pairwise meta-analyses.

*Methods:* Network meta-analyses can also be used to plan future trials. We introduce a four-steps framework to plan a new trial that aims to identify the optimal new design that will update the existing evidence to best serve timely clinical and public health decision making. The new trial designed within this framework does not need to include all competing interventions and comparisons of interest and can contribute direct and indirect evidence to the updated network meta-analysis. We present the method by virtually planning a new trial to compare biologics in rheumatoid arthritis and a new trial to compare two drugs for relapsing-remitting multiple sclerosis.

*Results:* A trial design based on updating the evidence from a network meta-analysis of relevant previous trials may require a considerably smaller sample size to reach the same conclusion compared with a trial designed and analyzed in isolation. Challenges in the approach include the complexity of the methodology and the need for a coherent network meta-analysis of previous trials with little heterogeneity.

*Conclusions:* When used judiciously, conditional trial design could significantly reduce waste in clinical research.

## BACKGROUND

The role of evidence synthesis in directing future research in general, and planning clinical trials in particular has been debated widely in the medical and statistical literature [1,2]. Researchers designing new studies possibly consult other relevant studies informally to inform their assumptions about the anticipated treatment effect, variability in the health outcome or response rate. Empirical evidence has shown however that most randomized controlled trials (RCTs) do not consider systematic reviews or meta-analyses to inform their design [3–7]. One potential reason is that the available conventional meta-analyses had addressed only a subset of the treatment comparisons of interest that were being considered for the new trial. It is possible that the treatment comparison of interest has not been examined before and consequently there is no prior direct evidence to inform the future trial. However, the treatments of interest might have been compared to a common reference treatment in previous studies yielding indirect evidence about the comparison of interest.

In recent years methods to synthesize evidence across trials of multiple comparisons, called network meta-analysis, have become more widely used [8]. In network meta-analyses direct evidence from trials comparing treatments of interest (for example treatments A and B) and indirect evidence from trials comparing the treatments of interest with a common comparator (for example A and C, and B and C) is synthesized. Methods to plan a clinical trial specifically to update a pairwise meta-analysis were proposed ten years ago [9–11] and recently have been extended to the setting of network meta-analysis [12,13].

For example, a network meta-analysis based on 26 trials showed in 2015 that percutaneous coronary intervention with everolimus-eluting stents (EES) is more effective in the treatment of in-stent restenosis than the widely used intervention with drug-coated balloons, based on direct and indirect evidence [14]. Already in 2013, evidence from 19 trials favored EES but the estimate was imprecise with wide confidence intervals and entirely based on indirect comparisons. Two additional trials of drug-eluting balloons versus EES were subsequently published, which included 489 patients with in-stent restenosis (see Appendix 1). The inclusion of these two trials in the network meta-analysis rendered the evidence for the superiority of EES conclusive. The two studies were planned independently of each other, and without taking the existing indirect evidence into account. In fact, a single study with 232 patients would have sufficed to render the estimate from the network meta-analysis conclusive. In other words, the evidence could have become available earlier, with fewer patients being exposed to an inferior treatment.

In this article we illustrate how future trials evaluating the efficacy or safety of interventions can be planned based on updating network meta-analysis of previous trials, using an approach we call *conditional trial design*. Conditional trial design aims to identify the optimal new trial conditional on the existing network of trials to best serve timely clinical and public health decision making. Based on recent methodological work from our group and others, we discuss advantages and disadvantages of the approach.

## METHODS

We first describe the conditional trial design and then we discuss important challenges that are likely to be encountered in application.

### Conditional trial design

The approach is a step-wise process outlined below.

*Identify or perform a coherent network meta-analysis that addresses your research question*

The starting point is a network meta-analysis about the relative efficacy or safety of competing interventions for the same health condition. The network meta-analysis should be the result of a methodologically robust systematic review that aims to reduce the risk of publication and other bias. We also assume that between-study heterogeneity is low to moderate and the assumption of coherence is justified. In other words, for each treatment comparison the results across the different studies are reasonably similar (low heterogeneity, Box 1) and direct and indirect evidence are in agreement (coherence). Evaluation of homogeneity and coherence in a network can be performed using appropriate statistical quantities and tests and by comparing the characteristics of the studies (Appendix 3).

*Define the targeted comparison or comparisons between the treatments of interest whose relative effects you want to measure.*

In a next step the comparison or comparisons of interest, i.e. the ‘*targeted comparisons*’ need to be defined. Of note, network meta-analyses typically include comparisons with placebo or with older treatments that are not of interest per se, but may provide indirect evidence on the targeted comparisons. Depending on the clinical context the process of defining the targeted comparisons may be complex, involve many stakeholders, and might result in several targeted treatment comparisons.

*Decide whether the network meta-analysis answers the research question*

For each targeted comparison we need to establish whether the available evidence from the network meta-analysis is conclusive or not and if not, that a further trial or further trials are indeed warranted. An estimate of a relative effect might be characterized as inconclusive if the confidence interval around it is wide and includes both a worthwhile beneficial and harmful effect and generally includes values that could lead to different clinical decisions. When the hierarchy of treatment effectiveness is of interest, we consider the available evidence as inconclusive if the ranking of the treatments is imprecise i.e. the probabilities of a treatment of being best or worst are similar.

*Estimate the features of a future study that will update the network to answer the research question*

Once it is clear that additional evidence is required, we can proceed to planning the next trial. The new trial could compare the treatments in the targeted comparison or other treatments included in the network (‘*tested comparison*’). In this case the trial will contribute indirect evidence to the targeted comparison via the updated network. The choice of the targeted comparison depends on the research priorities in the clinical field and the values of the stakeholders involved in planning the research agenda. In contrast, decisions about the comparison actually tested in the new trial involve primarily practical considerations related to the feasibility of the trial (e.g. recruitment rate with the various treatment options) and sample size. In this article we focus on the sample size criterion; we would choose to test the comparison which minimizes the required sample size, typically that is the targeted comparison.

We have previously proposed a statistical methodology to estimate the sample size for a new trial within a conditional framework that takes the available evidence into account (Appendix 2). This methodology has two variants. The first one is based on the classical hypothesis testing concept. The sample size is estimated as a function of the conditional power; that is the power of the updated network meta-analysis when a new study is added and uses extensions of classical power calculations [12]. The second variant is based on estimating treatment effects rather than testing. The sample size is calculated so as to improve the precision of the estimate from the updated network meta-analysis so that clinically important effects can either be confirmed or excluded [13]. For both approaches the required sample size will be lower for a conditionally designed trial compared to a trial designed and analyzed in isolation. The two methods are essentially equivalent, but practical considerations may lead to a preference of one over the other. For example, if the ranking of several treatments is of interest, then the targeted comparisons are all comparisons between them and sample size calculations are more easily done using the second variant.

If the tested comparison is not the same as the targeted comparison the study will improve the precision in the estimates of the targeted comparison by contributing to its indirect evidence, but it will require a larger sample size to achieve the same level of precision or power compared to a study of the targeted comparison. Indirect evidence may be preferable in some situations, for example if one of the treatments of interest is more invasive or more inconvenient than the comparator of interest. Careful inspection of the existing network graph is important if the new trial is to provide indirect evidence: links between the treatments tested and the treatments of interest are required. Note that equipoise between the tested treatments will always need to be documented.

### Limitations of the conditional planning framework

A number of factors will limit the pragmatic applicability of the conditional planning framework. For instance, it will be irrelevant to the planning of a trial to evaluate a new intervention, or when the primary outcome of interest has not been considered in previous trials. Network meta-analysis rests on the assumption of coherence and only under this condition can be considered as the basis for trial design. However, evaluation of the assumption is challenging in practice (see Appendix 3), particularly for poorly connected networks with few studies. We have also stressed that between-study heterogeneity should be low for the total network and the targeted comparison in particular. If heterogeneity is large, the contribution of a single study, even if the study is large, is low and the precision of the treatment effect estimated by the network meta-analysis remains low [9,10,12]. Indeed, heterogeneity sets an upper bound to the precision of the summary effect beyond which precision cannot be improved: the upper bound of the attainable precision is equal to the number of studies divided by the heterogeneity variance (see formula in Appendix 4). Additionally, as heterogeneity increases interpretation of the synthesis becomes more problematic.

Estimation of heterogeneity requires several studies to be available for the targeted comparison. In the absence of many direct studies, the common heterogeneity estimated from the network can be considered along with empirical evidence, specific to the outcome and treatment comparison (Box 1). If there are is substantial heterogeneity it will not be possible to design a single trial based on the conditional trial design approach. If heterogeneity is large, and the sources of heterogeneity are well understood, planning several smaller studies instead, for example in patients with different characteristics, may be a more powerful approach than planning a single large trial in one patient group (Box 1).

## APPLICATION

We considered two examples from the published literature. We first consider the case of biologic and conventional disease-modifying anti-rheumatic drugs (DMARDs) for rheumatoid arthritis (RA). A second example, of treatments for multiple sclerosis, is also presented.

### Treatments for Rheumatoid Arthritis

Methotrexate is the ‘anchor drug’ for RA and recommended as the first DMARD. It is unclear, however, whether a biologic DMARD should be added to the initial therapy or whether initial therapy should be based on the less costly triple therapy of conventional DMARDS, i.e. methotrexate plus sulfasalazine plus hydroxychloroquine.

A recent network meta-analysis of randomized controlled trials in methotrexate-naïve patients concluded that triple therapy and regimens combining biologic DMARDs with methotrexate were similarly effective in controlling disease activity (Figure 1) [15]. Heterogeneity was low to moderate; the between-study variance was τ^2^=0.03. The assumption of coherence was deemed plausible after considering trial characteristics, although the lack of direct evidence for many comparisons does not allow formal statistical evaluation.

**Figure 1.**
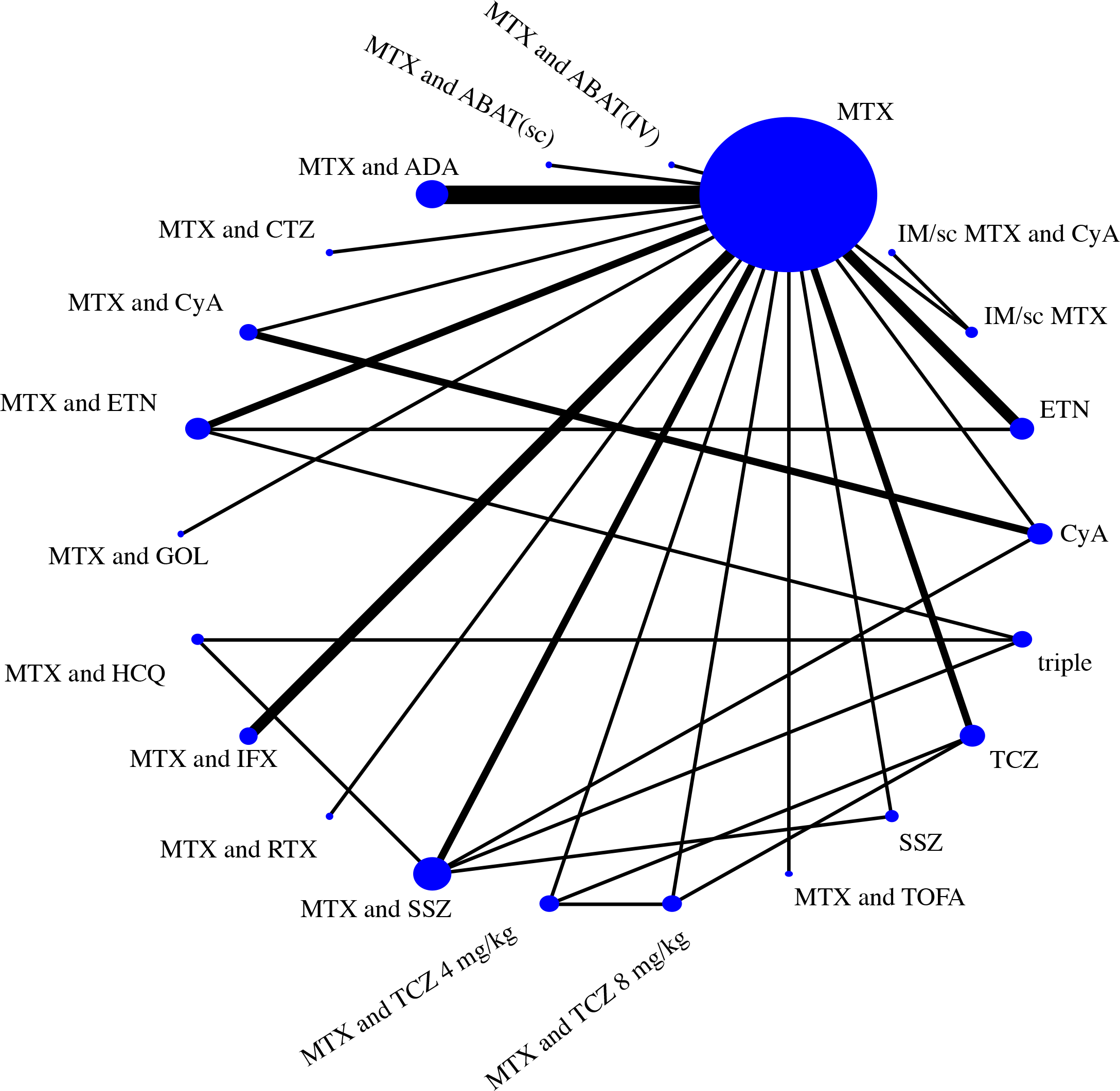
Network of evidence of treatments for rheumatoid arthritis. Adapted from [15]. ABAT=abatacept; ADA=adalimumab; AZA=azathioprine; CQ=chloroquine; ETN=etanercept; Ctl-certolizumab; CyA=cyclosporin; GOL=golimumab; HCQ=hydroxychloroquine; IFX=infliximab; IM=intramuscular; IR=inadequate response; iv=intravenous; LEF=leflunomide; MTX=methotrexate; RTX=rituximab; sc=subcutaneous; SSZ=sulfasalazine; TOFA=tofacitinib; TCZ=tocilizumab, triple= methotrexate plus sulfasalazine and hydroxychloroquine

Methothrexate combined with etanercept was associated with higher average response (defined as the American College of Rheumatology (ACR) 50 response) compared to all other treatments in the network. Synthesis of evidence from the one direct trial (with 376 patients [16]) with indirect evidence in the network favored methotrexate plus etanercept over the triple therapy in terms of efficacy but with considerable uncertainty: the odds of response were lower with triple therapy (odds ratio 0.71) but the 95% confidence interval was wide (0.42 to 1.21). Clearly, the available evidence on the targeted comparison is inconclusive. A more precise estimate of the comparative efficacy of the two therapies would add clarity to whether any added benefit of etanercept justifies the extra cost and thus would be of interest to guideline developers and reimbursement agencies.

A trial directly comparing methotrexate plus etanercept and triple therapy is the most efficient approach to generating the required additional evidence. Figure 2 shows the gain in power for a new etanercept and methotrexate vs triple conventional DMARD therapy trial when designed conditional on the existing network of trials. Assuming equal arm allocation at randomization, a trial with 280 patients in total will enable the updated network meta-analysis to detect an odds ratio of 0.71 with 80% power. The corresponding sample sizes for a conventional pairwise meta-analysis (pooling with the existing study [16]) or a standalone trial are 790 and 1084 patients, respectively.

**Figure 2.**
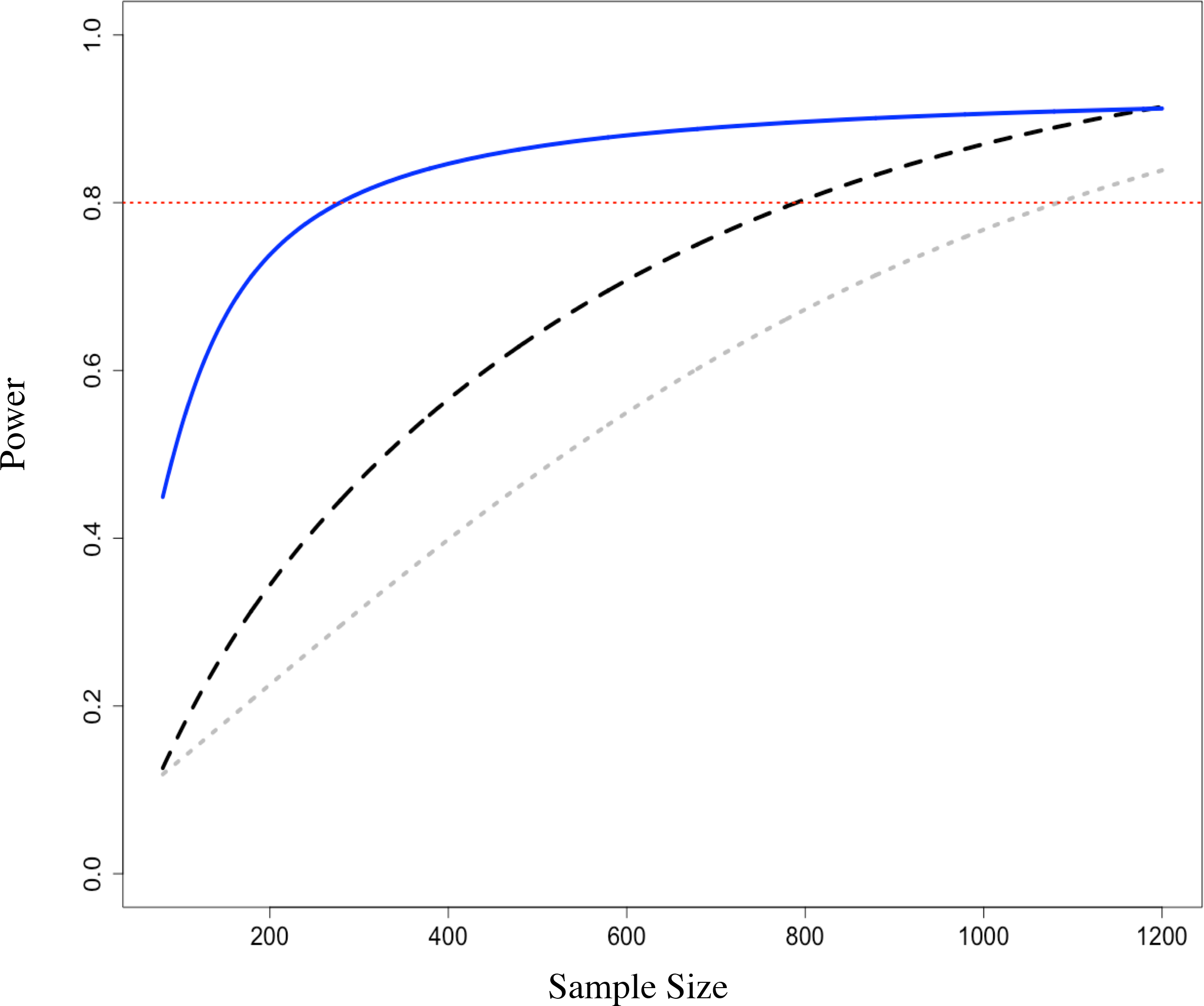
Power and conditional power to detect a difference in response between *triple therapy versus methotrexate combined with etanercept* as a function of the sample size. The difference anticipated in a new study was of OR=0.71 in American College of Rheumatology (ACR) 50 response favouring methothrexate/etanercept. Power for a single randomised trial considered in isolation (dotted line), conditional power for a fixed effect pairwise meta-analysis (dashed line, two studies) and network meta-analysis (solid line). Expected event rate in the triple therapy was assumed equal to the average observed in the network (49%). Calculations were performed using the conditional power method [12].

Appendix 2 presents technical guidance to allow inclined readers to re-produce results.

### Treatments for relapsing-remitting multiple sclerosis

In 2015, the association of British Neurologists published guidelines about choosing treatments for multiple sclerosis (MS) [17]. The guidelines classify fingolimod, dimethyl fumarate, beta-interferon and glatiramer acetate in the category of moderate efficacy. They recommend that patients with relapsing-remitting condition start with fingolimod or dimethyl fumarate because they are most effective and are administered orally.

A relevant NMA with low heterogeneity (τ^2^=0.01) and no serious concerns about incoherence was published in the same year (Figure 3) [18]. The results, while broadly in line with [17], show some differences between the drugs in the moderate efficacy group. Fingolimod is shown to considerably decrease the rate of relapses compared to dimethyl fumarate while evidence is weak to support the advantage of fingolimod over glatiramer acetate (Risk Ratio RR 0.88 (0.74, 1.04)). As reported in the guidelines, glatiramer acetate has been used “extensively for decades in MS” and its safety profile is well understood, in contrast to fingolimod, which is a newer agent (only two placebo-controlled trials were included in the review). Patients, their doctors and guideline developers might want stronger evidence against the similar efficacy of these two interventions.

**Figure 3.**
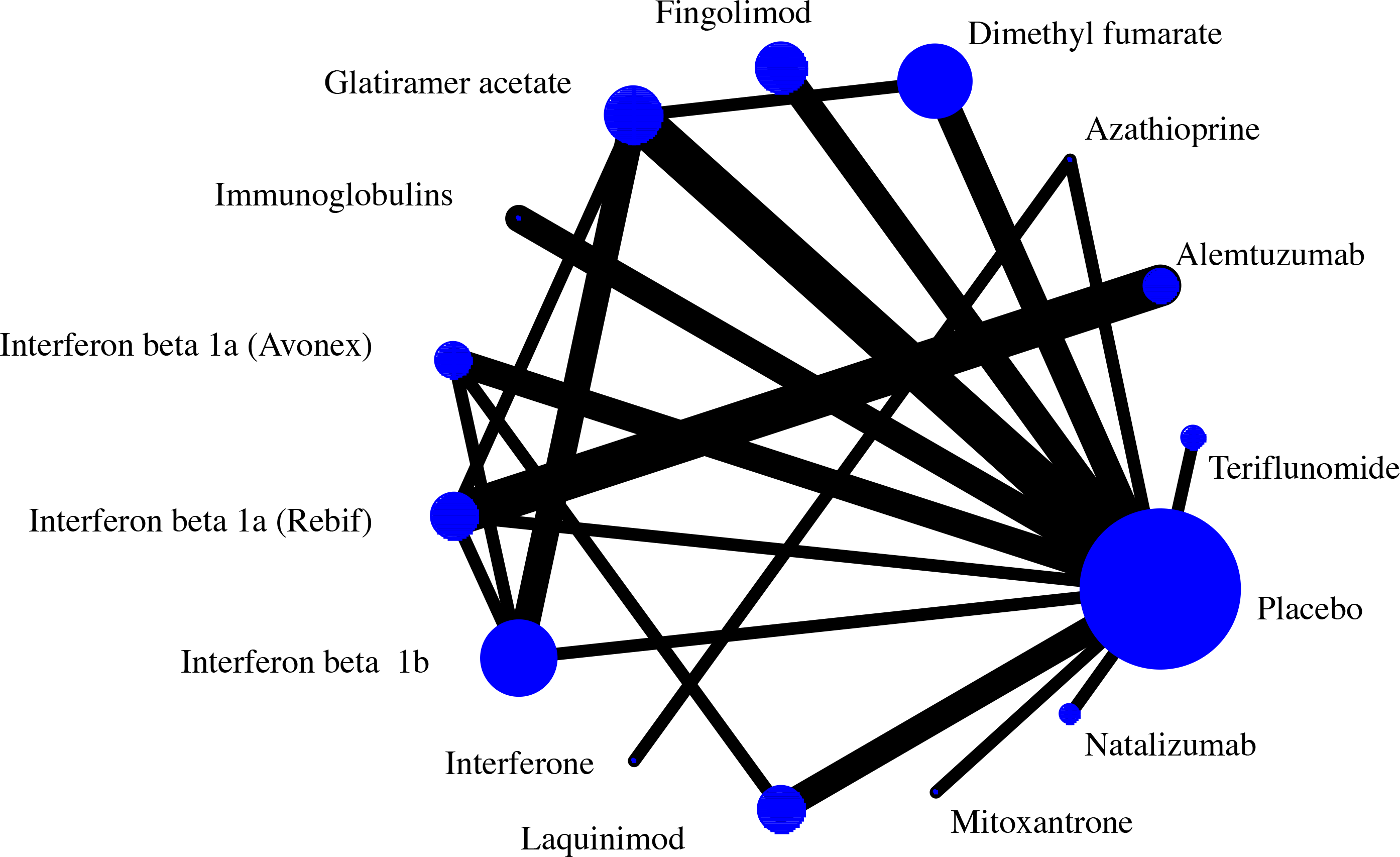
Network of evidence of treatments for multiple sclerosis. Adapted from [18].

We estimated the sample size required in a new trial that compares fingolimod and glatiramer acetate to update the network using the precision-minimization approach. Assuming a 51% relapse rate for glatiramer acetate 552 participants are needed in a fingolimod vs glatiramer acetate study to update the NMA and exclude a risk ratio of one in the fingolimod vs glatiramer acetate risk ratio. An independently designed and analysed study would need 1200 participants to achieve the same level of precision. Note that the targeted comparison does not have direct evidence and fingolimod is connected only to placebo; consequently, precision can be improved only if fingolimod is included in the new study. A fingolimod versus dimethyl fumarate study would need a much larger sample size to achieve the same level of precision in the fingolimod vs glatiramer acetate updated RR (2742 participants).

## DISCUSSION

We present a framework of conditional trial design that has the potential to reduce the resources needed to answer a clinical question of relevance to health-policy. Reduced sample size and flexibility in the randomized arms included are the main advantages of the method. However, it is only applicable to cases where a well-conducted NMA is available (or is possible to undertake) and the underlying treatment effect under investigation is not expected to have important variability across trial settings.

Efficient trial design is paramount to both public and private funders of research: investing in a clinical trial consumes funds that could instead be diverted to other research activities. The proposed conditional trial design has the potential to bring substantial benefits to drug licensing agencies and health technology assessment (HTA) bodies. The approach depends on the availability of a coherent and fairly homogeneous network of trials, relevant to the research question the future trial aims to answer. Consequently, the feasibility and the added benefit (in terms of reduction in sample size) of the conditional trial design remain to be empirically tested. User-friendly software to simplify the process will be required and trialists will need to familiarize themselves with the technique. More importantly, wide adoption of the conditional trial design will require a shift in the current paradigm among regulators, reimbursement decision bodies and funders of research.

In recent years, drug licensing agencies such as the Food and Drug Administration (FDA) and European Medicines Agency (EMA) have embraced conditional, restricted approval mechanisms instead of making an initial binary decision as to whether a new treatment should be approved or rejected. Following initial market entry, in many health care systems, HTA bodies are tasked with advising practice guideline development panels and payers about the therapeutic and economic value of treatments. Similar to drug licensing agencies, HTA bodies increasingly require the generation of additional evidence under so-called conditional coverage options [19]. For example, UK’s National Institute for Health and Care Excellence can restrict the use of a new treatment to research participants in its “only in research” designation. In this capacity, drug licensing agencies and HTA bodies are well positioned to strengthen the link between existing and future research, and ensure that design features of future clinical trials are informed by the totality of available relevant evidence. A research agenda of national and international public and private funders of trials which is streamlined with the evidence needs of regulatory agencies, HTA bodies and guideline developers could benefit from the conditional planning of trials and generate meaningful and relevant evidence faster and more efficiently.

## CONCLUSION

The role of network meta-analysis in guideline development, health technology assessment [19] and drug licensing [20] is increasingly recognized [21]. Conditional trial design extends the use of network meta-analysis to the efficient design of future trials.

As Altman pointed out in 1994, we need less but better research, and research done for the right reasons [22]. Since then, specific recommendations have been made to increase the value of clinical research, and reduce waste [23]. We believe that conditional trial design, used judiciously based on a homogenous and coherent network of the available controlled trials, can obtain conclusive results earlier, facilitate timely decision making and reduce research waste.

### Ethics approval, consent to participate

Not required

### Consent for publication

Not required

### Availability of data and material

The datasets used in the examples are publicly available in the original reviews and are also presented in the Appendix. The RA example (data and analysis) is also available in the GitHub repository.

## Acknowledgements

We would like to thank Glen Hazlewood for helpful clarifications regarding the data and analysis of the RA dataset.

## Authors’ contributions

GS and ME conceived and designed the study. AN assisted in the design of the study and performed the main analyses. HN provided input about the implications for policy, AS and ST commented on and made suggestions for the methodology for sample size calculations and SR participated in the analysis of the RA example. All authors critically revised the manuscript, interpreted the results and performed a critical review of the manuscript for intellectual content. GS, AN and ME produced the final version of the submitted article and all co-authors approved it. AN and GS had full access to all data in the study and take responsibility for the integrity of the data and the accuracy of the analysis. GS and EM are the guarantors.

The lead author (GS) affirms that the manuscript is an honest, accurate, and transparent account of the study being reported; that no important aspects of the study have been omitted; and that any discrepancies from the study as planned (and, if relevant, registered) have been explained.

## Box 1 A suggested strategy for measuring and addressing heterogeneity when planning a future study to update a network meta-analysis

1. *Measure and characterize heterogeneity.* Latest methodological developments encourage meta-analysts to estimate the heterogeneity variance which measures the variability of the true treatment effect and compare it to its expected value. Within the context of planning new studies, existing heterogeneity is large when even a new study with several thousands of participants fails to produce effect sizes that are precise enough to enable decision making. Consequently, characterizing heterogeneity as large or small depends on the context. We recommend that when the heterogeneity variance is below the expected, sample size calculations are carried out; if the estimated sample size is unrealistically large, then heterogeneity is clearly too large to proceed with conditional trial design.
2. *Understand heterogeneity.* Identify potential effect modifiers and explore the changes of the treatment effect in various *subgroup analyses* using study-level and patient-level characteristics (such as risk of bias, trial setting). The impact of continuous characteristics (such as trial duration or sample size) can be explored using *meta-regression models.* Interpret with caution any findings from patient-level covariates as they can be subject to ecological bias.
3. *Reduce heterogeneity.* If some variables are associated with heterogeneity, then consider the treatment effects about a particular group of patients and trial settings. For example, investigators might choose to restrict the analysis in low risk of bias or recent studies, or estimate the treatment effect for a very large study (from a meta-regression on sample size). Any posthoc decisions about the analyzed dataset need to be clearly documented to avoid selection bias.
4. *What to do in the presence of large, unexplained heterogeneity*. If the heterogeneity remains substantial even after efforts to pinpoint its source and reduce it, then investigators have the following options a) collect individual-patient data to better investigate the role of patient characteristics in modifying the treatment effect and go to step 2 above b) develop assumptions about what is possibly causing the heterogeneity and plan a new exploratory study to investigate these assumptions. For example, if investigators believe that treatment dose is associated with differential effects (and this information is scarcely reported in the included studies) a new multi-arm study can be planned to investigate this assumption.

